# A Third-generation Lentiviral Vector System for Flow Virometry-based Analysis of HIV-1 Env and Other Viral Envelope Proteins

**DOI:** 10.64898/2026.10.11.755930

**Authors:** Paula I. Busskamp Castro, Janine Parreuter, Charlotte Brands, Ana-Maria Perkovic, Benton van Bilsen, Bernd Giebel, Jens Bohne, Tobias Tertel, Jamal Ghanam, Basant K. Thakur, Christina B. Karsten

## Abstract

Individual human immunodeficiency virus (HIV) particles can differ in their envelope protein (Env) despite being genetically identical. This non-genetic diversity can influence antibody recognition and has been implicated in the formation of persistent virus fractions that escape complete inhibition by broadly neutralizing antibodies. Investigating non-genetic diversity in HIV and other viruses requires approaches capable of characterizing antibody binding to individual viral particles. Flow virometry provides a promising tool for single-particle analysis, but a system combining high biosafety with specific particle labeling and versatility is lacking. Here, we developed a third-generation lentiviral vector system specifically designed for flow virometry.

Fluorescent proteins were incorporated into the HIV-1 Gag polyprotein between the matrix- and capsid-encoding regions, flanked by viral protease cleavage sites, creating the packaging constructs pLentiFlow-eGFP and pLentiFlow-mCherry. Lentiviral particles (LVPs) based on pLentiFlow vectors were fluorescent and retained infectivity comparable to LVPs generated with the parental vector system despite reduced particle production. We optimized LVP detection on a CytoFLEX nano and defined a linear p24-based concentration range for reliable single-particle analysis. Using complementary controls, we demonstrated that background from the assessed sources accounted for no more than approximately 3% of events within a broad eGFP+ gate and confirmed that the detected fluorescent particles were predominantly membrane-enclosed. As a proof of concept, LVPs were pseudotyped with vesicular stomatitis virus glycoprotein, severe acute respiratory syndrome coronavirus 2 spike, or HIV-1 Env. Envelope-specific antibody binding was detected using 8G5F11, CR3022 or PGT128, respectively, demonstrating that different viral surface proteins can be analyzed on a common particle backbone.

Together, these results establish a third-generation lentiviral vector system for flow virometry that combines intrinsic particle labeling with high biosafety and flexible pseudotyping. The system provides a platform for investigating envelope protein-associated non-genetic diversity and its consequences for antibody recognition in HIV and other viruses.

**Author summary:** Recent studies suggest that non-genetic diversity in the envelope protein of the human immunodeficiency virus (HIV) influences antibody recognition, a phenomenon potentially relevant to other viruses as well. Understanding such non-genetic differences may aid the development of antibody-based vaccines and therapies. To assess envelope protein-associated non-genetic diversity within viral swarms, methods are needed to analyze individual viruses. Flow virometry is a promising approach but current methods face challenges in combining single-particle detection by flow cytometry with high biosafety and versatility.

Here, we present a third-generation lentiviral vector system that generates fluorescent lentiviral particles (LVPs) for direct detection. These LVPs are produced from a reduced and split HIV genome and cannot produce infectious virus after infecting a cell, providing a high level of biosafety. The incorporated viral envelope proteins can be exchanged, allowing the study of HIV and other viruses. We characterize the LVPs, optimize their detection, establish conditions for reliable analysis of individual particles, and demonstrate the versatility of the system by detecting antibody binding to envelope proteins from HIV and two other viruses. By combining biosafety and versatility, this system broadens access to flow virometry and enables the study of non-genetic differences within swarms of HIV and other viruses.

## Introduction

Genetic diversity of the human immunodeficiency virus (HIV) represents a major barrier to vaccine development [1] and underlies the need for combination antiretroviral therapy to control established infection [2, 3]. However, comparative studies investigating genetically identical HIV or its model simian immunodeficiency virus (SIV) produced by CD4+ T cells and macrophages have revealed producer cell-dependent differences in virion composition and phenotype that arise independently of viral genetic variation [4–12], demonstrating that genetically identical viral populations can acquire distinct properties depending on their cellular origin. Importantly, non-genetic diversity also occurs between individual particles within genetically identical viral populations, with important consequences for antibody recognition. In particular, genetically identical HIV-1 or HIV-1 Env-bearing lentiviral particle (LVP) populations that are generally sensitive to a given broadly neutralizing antibody (bnAb) can contain persistent fractions that remain infectious in its presence, a phenomenon observed across bnAbs targeting distinct Env epitopes [13]. Further, in a rhesus macaque simian-HIV challenge model, a larger persistent fraction was associated with reduced sterilizing protection following bnAb administration, although the extent to which differences in bnAb potency contributed to this outcome remains unclear [14]. Several Env-associated features have been implicated as cause for the persistent fraction, with heterogeneity in Env glycosylation [15, 16], stability [17], and conformation [18] contributing to differential antibody susceptibility among genetically identical particles. Interest in persistent fractions during antibody neutralization extends beyond HIV, with both persistent fractions and the effects of variation in antibody-targeted viral surface proteins on antibody recognition investigated in several human pathogenic viruses, including poliovirus, severe acute respiratory syndrome coronavirus 2 (SARS-CoV-2), West Nile virus, and herpesviruses, among others [19–24]. Together, the studies on HIV and other viruses have provided valuable insights but also highlighted that the formation of a persistent fraction is likely complex and multifactorial, with the underlying mechanisms potentially varying between individual envelope protein-antibody interactions. Thus, further investigation is needed to comprehensively understand this phenomenon and its implications for antibody-based vaccines and therapies.

Detailed characterization of non-genetic variation in HIV-1 Env and other viral envelope proteins requires efficient approaches capable of assessing antibody binding at the level of individual viral particles. Flow virometry, the flow cytometric analysis of viral particles, is an emerging technology that enables the characterization of viral subpopulations and the isolation of defined particle populations for downstream analyses. However, the small size of viral particles leads to minimal light scattering. More importantly, their discrimination is complicated by extracellular vesicles (EVs) present in virus stocks as well as similarly sized particles that may be present in reagents and buffers and exhibit overlapping light-scattering characteristics [25]. Several strategies have therefore been developed to facilitate viral particle detection. Coupling viruses to larger beads increases their effective size, facilitating their visualization, but multiple viral particles may associate with a single bead, limiting true single-particle analysis. Antibody-based capture may additionally bias the analyzed viral population toward particles displaying the targeted antigen or epitope [26]. Alternatively, viral particles can be labeled with lipophilic or cytosolic dyes, but these dyes also label EVs of similar size present in virus preparations and therefore do not specifically identify viral particles [27].

More specific labeling for single-virus analysis can be achieved by directly labeling viral components. For HIV, incorporation of fluorescent proteins into Gag provides a particularly attractive strategy because it intrinsically labels viral particles independently of Env and therefore permits subsequent characterization of Env without using it for particle identification. The feasibility of this approach was demonstrated using the first-generation lentiviral vector HIV Gag-iGFP ΔEnv, which enabled direct visualization and sorting of intrinsically fluorescent LVPs by flow virometry [28, 29]. However, this system remains subject to substantial biosafety restrictions, limiting its accessibility for flow virometry applications. Depending on national regulations, work with genetically modified HIV and other viruses may require high-containment laboratories, which less commonly house suitable flow cytometers or cell sorters. Beyond regulatory requirements, reducing the biological risk associated with viral particle production and flow cytometric analysis is desirable, particularly for procedures carrying a potential risk for aerosol formation, such as flow virometric visualization and sorting [30, 31]. A further limitation is that establishing reliable flow virometry detection requires considerable optimization [26].

Using an otherwise consistent particle system with interchangeable envelope proteins therefore avoids the need to establish and optimize a separate particle system for each virus of interest. Such a system would permit comparative analyses of different Env variants or envelope proteins from different viruses while maintaining a common fluorescent particle backbone and established acquisition conditions.

In summary, there is a need for a system combining bead-independent, specific particle labeling with high biosafety and flexible pseudotyping to facilitate the investigation of non-genetic envelope-associated diversity in HIV and other viruses. In the present study, we therefore developed a third-generation lentiviral vector system for flow virometry designed to address these requirements.

## Material and Methods

### Plasmids

The parental third-generation lentiviral vector system consisted of the packaging plasmid pMDLg/pRRE, encoding HIV-1 Gag and Pol, and the Rev expression plasmid pRSV-Rev [32]. For pseudotyping, the envelope expression plasmids pMD2.G (vesicular stomatitis Indiana virus glycoprotein (VSV-G)), pDOLHIVenv (HIV-1 NL4-3 Env), or pCAGGS-FL CoV-2 Spike (SARS-CoV-2 Wuhan-Hu-1 spike) (all BEI Resources) were used.

The fluorescent packaging vectors pLentiFlow-eGFP and -mCherry were generated as follows. First, the sequence of pMDLg/pRRE was PCR-amplified using primers annealing to the 5′ and 3′ ends of the HIV-1 protease cleavage site (SQNYPIVQ) located between the matrix and capsid coding sequences, linearizing the backbone and duplicating the cleavage site. Next, eGFP and mCherry were PCR-amplified from pMA-RQ-eGFP and pcDNA-mCherry (both this study), with primers introducing the overlaps required for DNA assembly. Following DpnI digestion and gel purification of all PCR products, the fluorophore coding sequences were inserted between the duplicated HIV-1 protease cleavage sites using the NEBuilder HiFi DNA Assembly Master Mix according to the manufacturer’s instructions. The resulting plasmids were verified by Sanger sequencing.

The transfer plasmid pRRL.SIN.cPPT.EF1α-TagBFP.WPRE was derived from pRRL.SIN.cPPT.SF-GFP.WPRE [33]. To enable directional cloning, one of the two BamHI and NotI restriction sites was removed by site-directed mutagenesis. The spleen focus-forming virus promoter was then replaced with the human elongation factor 1 alpha (EF1α)-promoter of pSBbi-RP [34] using BamHI/NheI. TagBFP was subsequently inserted using NotI/BamHI from pcDNA-TagBFP (this study). All newly generated plasmids were verified by sequencing.

For recombinant antibody production, the plasmid pair pFUSEss-CHIg-hG1-CR3022 heavy chain and pFUSEss-CLIg-hκ-CR3022 light chain (BEI Resources) was used to express the SARS-CoV-1/2-specific antibody CR3022.

### Cell culture

HEK293T and TZM-bl cells were maintained in complete DMEM (Dulbecco’s Modified Eagle Medium (Gibco) supplemented with 10% fetal bovine serum (FBS) and 2 mM L-glutamine (both Capricorn Scientific)) with 10^5^ units/L penicillin, and 100 mg/L streptomycin (both Capricorn Scientific) at 37°C and 5% CO₂. FreeStyle 293-F cells were cultured in FreeStyle 293 Expression Medium (Gibco) at 37°C, 8% CO₂, 70% humidity and 135 rpm.

### Production of antibodies

The human monoclonal antibody CR3022 against SARS-CoV-1/2 spike was produced by transient transfection of FreeStyle 293-F cells using polyethylenimine (PEI; 1 mg/mL; Kyfora Bio). For antibody production, FreeStyle 293-F cells were transfected at a density of 1.2 × 10⁶ cells/mL. Heavy- and light-chain expression plasmids were mixed at equal ratios using a total of 250 µg plasmid DNA per 500 mL culture. DNA and 750 µL PEI were diluted separately in 12.5 mL Opti-MEM (Thermo Fisher Scientific), sterile-filtered, combined after 5 min incubation at room temperature, and incubated for an additional 30 min to allow complex formation. The transfection mixture was then added dropwise to the cells.

Five days after transfection of FreeStyle 293-F cells, supernatants were harvested, centrifuged at 3,500 × g for 20 min, and filtered using 0.8/0.2 µm VacuCap filtration devices (Pall). Antibodies were purified by affinity chromatography using HiTrap Protein G columns with an ÄKTA pure 25L FPLC system (both Cytiva), concentrated using Amicon Ultra centrifugal filters with a 3 kDa molecular weight cut-off (Merck), and further purified by size-exclusion chromatography using a Superdex 200 Increase 10/300 GL column. Protein concentrations were determined using a BioSpectrometer (Eppendorf).

### Production of pseudotyped LVPs

HEK293T cells were seeded one day before transfection at 2.2 × 10⁶ cells per T25 flask or 6.6 × 10⁶ cells per T75 flask in complete DMEM. On the day of transfection, the medium was exchanged 2 h prior to transfection. For transfection, a total of 2,500 fmol (T25) or 7,000 fmol (T75) plasmid DNA was used. Packaging plasmid, Rev expression plasmid, envelope expression plasmid, and transfer plasmid were mixed at a molar ratio of 20:15:18:36. The DNA mixture was diluted in 220 µL (T25) or 660 µL (T75) PBS. PEI was diluted separately in the same volume of PBS at a 3:1 PEI:DNA mass ratio. After 10 min of incubation, the diluted DNA and PEI solutions were combined and incubated for 30 min at room temperature to allow complex formation. The resulting DNA/PEI complexes were added dropwise to the cells, with transfection conditions adjusted to maintain a final PEI concentration below 4.5 µg/mL in the culture medium.

After overnight incubation at 37°C and 5% CO₂, the medium was replaced with complete DMEM. Two to three days after transfection, cell culture supernatants were harvested, centrifuged at 2,300 × g for 5 min, filtered through 0.45 µm PES membrane syringe filters (TPP), aliquoted, and stored at −80°C until use.

### Quantification of p24 by enzyme-linked immunosorbent assay (ELISA)

The p24 content of LVP preparations was quantified using the HIV-1 p24CA Antigen Capture Assay Kit (Frederick National Laboratory for Cancer Research) according to the manufacturer’s instructions. Absorbance was measured at 450 nm with a reference wavelength of 570 nm using a Spark microplate reader (Tecan).

### Infectivity assays

For infectivity assays, TZM-bl cells were seeded in 12-well plates at 2.2 × 10⁵ cells per well in 1 mL complete DMEM supplemented with 25 mM HEPES (Gibco). Cells were incubated with 220 µL of diluted LVP preparations in the presence of 20 µg/mL DEAE-dextran (Sigma-Aldrich) in a final volume of 2 mL per well. After 2–3 days of incubation at 37°C and 5% CO₂, cells were harvested by trypsinization using 0.05% trypsin/0.02% EDTA in PBS (PAN Biotech), washed with PBS (400 × g for 5 min), and stained with the LIVE/DEAD Fixable Near-IR Dead Cell Stain Kit (Thermo Fisher Scientific) according to the manufacturer’s instructions. Cells were washed again and fixed with 300 µL of 4% paraformaldehyde (PFA; RotiHistofix Formaldehyde, Carl Roth) for 15 min at room temperature protected from light, and washed once more. Before acquisition on a FACSymphony A5 flow cytometer (BD Biosciences), samples were filtered through cell strainers (35 µm, Corning), and the percentage of live TagBFP-positive cells was determined as a readout of LVP-mediated transduction. Data were analyzed with FlowJo software (version 10.10.0; BD Biosciences).

### Imaging flow cytometry

To determine LVP fluorescence by imaging flow cytometry, 2.2 × 10^5^ TZM-bl cells were incubated with LVPs pseudotyped with VSV-G (5 ng p24 per 1 × 10⁴ cells) in the presence of 20 µg/mL DEAE-dextran in complete DMEM in a total volume of 2 mL. Samples were incubated for 30 min at 4°C to allow particle attachment while preventing viral uptake. Subsequently, cells with attached LVPs were washed with PBS (5 min, 200 × g) and fixed with 4% PFA in PBS for 2 h at room temperature protected from light. Fixed cells were washed, stained with DAPI (Invitrogen) for 30 min, and washed again before acquisition.

Samples were acquired using an Amnis ImageStreamX Mk II imaging flow cytometer (Cytek Biosciences) equipped with an autosampler for U-bottom plates. Instrument settings were established using unstained cells, DAPI-stained cells without LVPs, and DAPI-stained cells with pLentiFlow-eGFP- or pLentiFlow-mCherry-based LVPs as fluorescence controls. Samples were measured for 20 min at 40× magnification with a medium flow rate of 2.5 µL/min. For detection, the following channels were used: brightfield (Ch01), eGFP (Ch02), mCherry (Ch03), and DAPI (Ch07). Data were analyzed using IDEAS software (version 6.3; Amnis/Cytek).

### Flow virometry

Flow virometry measurements were performed using a CytoFLEX nano flow cytometer (Beckman Coulter) in standard configuration. Prior to each experiment, CytoFLEX nano Daily QC Scatterspheres and Fluorospheres (both Beckman Coulter) were acquired according to the manufacturer’s instructions to verify instrument performance. For antibody staining of LVPs, in-house-produced CR3022, mouse anti-VSV-G (clone 8G5F11, Merck), and human anti-HIV-1 gp120 (PGT128, BEI Resources) were directly conjugated to the allophycocyanin (APC) using Lightning-Link conjugation kits (Abcam) according to the manufacturer’s instructions.

All sample and antibody dilutions were prepared in freshly filtered TNE buffer (10 mM Tris-HCl, pH 7.4, 150 mM NaCl, 2 mM EDTA; 0.05 µm-filtered). Prior to dilution, antibodies were centrifuged at 16,100 × g for 5 min at 4°C. Staining was performed in 96-well U-bottom plates in a total volume of 60 µL. The indicated concentrations of directly labeled antibodies were combined with LVPs equivalent to 1 ng p24 and mixed by pipetting. Following incubation for 1 h at room temperature in the dark, LVPs were pelleted by centrifugation at 16,100 × g for 20 min at 4°C and resuspended in 120 µL TNE buffer. Samples were subsequently adjusted to a final volume of 200 µL using 0.05 µm-filtered PFA prepared in TNE buffer to achieve a final PFA concentration of 4%.

Based on the initial LVP input of 1 ng p24, this corresponded to a nominal LVP concentration of 5 ng p24/mL. Samples were fixed for 10 h at 4°C. Fixed samples were stored at 4°C until acquisition. Standard controls included, as appropriate, TNE buffer only, individual antibodies diluted in TNE buffer, LVPs of the parental system, unstained LVPs, Triton X-100-treated LVPs, single-color-stained LVPs, and fluorescence-minus-one controls.

Unless otherwise indicated, samples were acquired for 60 s at a flow rate of 1 µL/min using a violet side scatter (VSSC)-1 gain of 1,250, a VSSC-2 gain of 500, and a VSSC-1-H triggering threshold of 2,500. For calibration of scatter and fluorescence measurements, Nanobead NIST Traceable Particle Size Standards (Polysciences) and ViroCheck NanoParticle Reference Kit (Invitrogen), respectively, were acquired alongside experimental samples. Data were acquired using CytExpert nano (version 1.2.0.34; Beckman Coulter). Using FCM_PASS_ (version 5.0.16; [35]), arbitrary VSSC-2 scatter signals were converted into estimated particle diameters based on a refractive index of 1.42 for enveloped viruses [36], and fluorescence signals were converted into equivalent reference fluorophore (ERF) units. Converted data were subsequently analyzed using FlowJo.

### Data management

The complete sequences of the newly generated plasmids pLentiFlow-eGFP, pLentiFlow-mCherry and pRRL.SIN.cPPT.EF1α-TagBFP.WPRE were deposited in GenBank (accession numbers: X–X). Supporting datasets were deposited in Dryad (Link), including raw flow cytometry and flow virometry data together with detailed experimental metadata according to the MIFlowCyt [37] and MIFlowCyt-EV [38] reporting frameworks.

## Results

### Engineering of the new third-generation lentiviral vector system for flow virometry

To combine targeted fluorescent particle labeling with an improved biosafety profile, we transferred this design concept to a well-established third-generation lentiviral vector system [32]. In this system, only three HIV-1 genes (*gag*, *pol*, and *rev*) are retained and distributed across the packaging plasmid pMDLg/pRRE, encoding Gag and Pol, and the Rev-expressing plasmid pRSV-Rev. The resulting LVPs are single-round infectious and can be pseudotyped with co-expressed envelope proteins of choice.

To generate a system suitable for flow virometry (Fig 1), the HIV-1 protease cleavage site (SQNYPIVQ) between matrix and capsid within pMDLg/pRRE was duplicated to create an insertion site for fluorescent proteins flanked by viral protease recognition sequences. The sequences of eGFP or mCherry were subsequently introduced between the duplicated cleavage sites using DNA assembly, generating the novel packaging constructs pLentiFlow-eGFP and pLentiFlow-mCherry. This design aimed to allow expression of Gag-fluorophore fusion proteins while maintaining the proteolytic processing sites required for fluorophore release during particle maturation.

**Fig 1.**
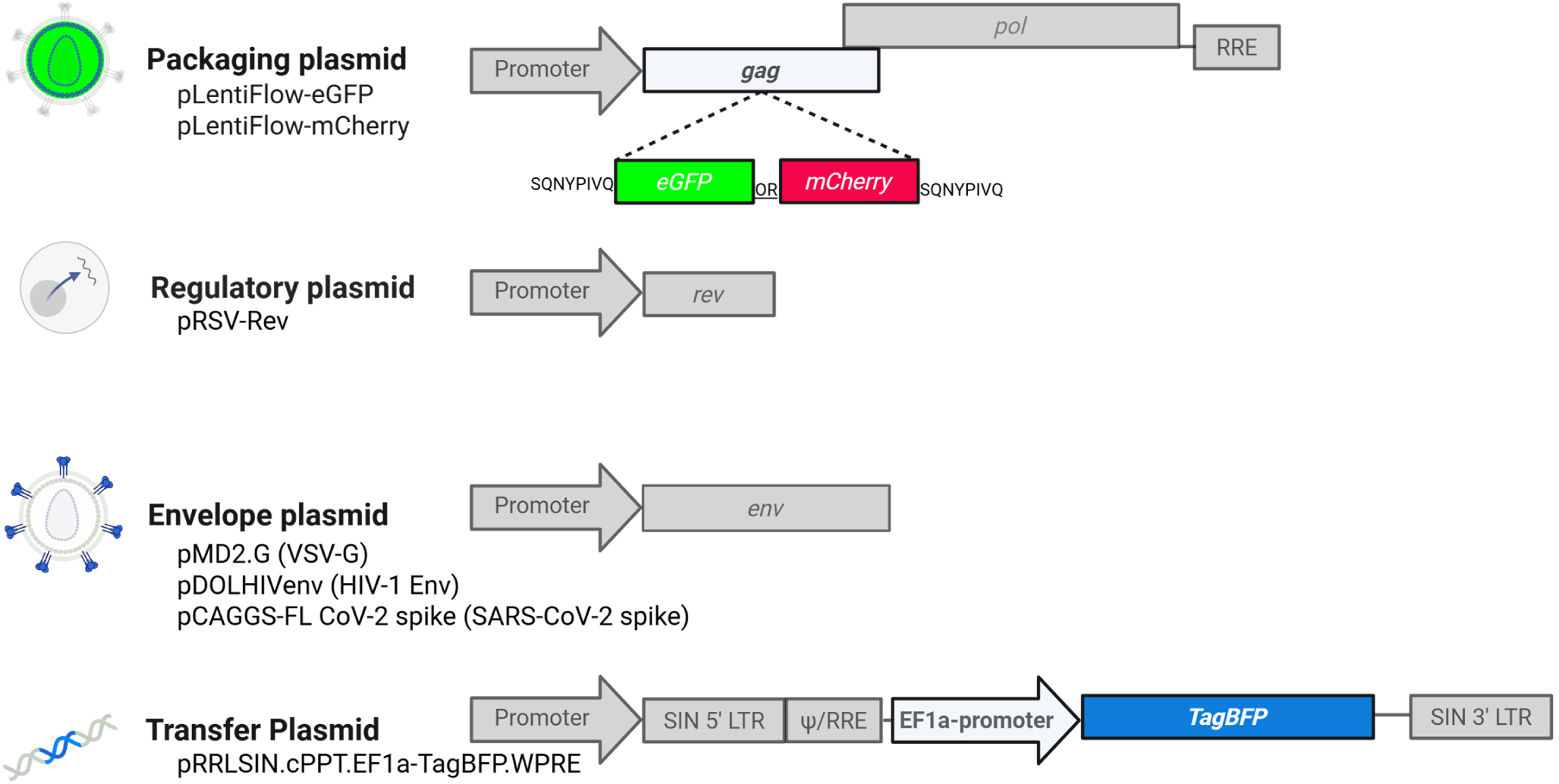
Schematic representation of the new third-generation lentiviral vector system for flow virometry. The parental third-generation lentiviral vector system is shown together with the modifications introduced for flow virometry. The respective fluorophore was inserted into the parental packaging plasmid pMDLg/pRRE between duplicated HIV-1 protease cleavage sites (SQNYPIVQ), resulting in pLentiFlow-eGFP and pLentiFlow-mCherry. The transfer plasmid was modified to encode TagBFP under the control of the human elongation factor-1 alpha (EF1α) promoter. Abbreviations: HIV-1 Env, human immunodeficiency virus-1 envelope protein; Ψ, RNA packaging signal; SARS-CoV-2, severe acute respiratory syndrome coronavirus-2; SIN 5′/3′ LTR, self-inactivating 5′/3′ long terminal repeat; RRE, Rev response element; VSV-G, vesicular stomatitis virus glycoprotein.

In addition, pRRL.SIN.cPPT.SF-GFP.WPRE [33], in which the long terminal repeat (LTR) becomes transcriptionally inactive following integration, was selected as transfer vector template. Using conventional restriction enzyme-based cloning, the original reporter cassette was modified to encode TagBFP under the control of the EF1α promoter, resulting in the transfer plasmid pRRL.SIN.cPPT.EF1α-TagBFP.WPRE.

The transfer plasmid pRRL.SIN.cPPT.EF1α-TagBFP.WPRE was designed to provide a spectrally distinct genetic reporter from the fluorescently labeled Gag proteins incorporated into pLentiFlow-derived LVPs, enabling the independent assessment of particle-associated fluorescence and successful transduction in subsequent experiments. At the same time, the use of a SIN transfer vector maintains the improved biosafety profile of the third-generation lentiviral vector system, including lowering the likelihood of recombination events with endogenous retroviral elements and reducing the risk of insertional oncogenesis [39, 40].

For all following analyses, LVPs were produced by co-transfection of pLentiFlow-eGFP or pLentiFlow-mCherry together with pRSV-Rev, pRRL.SIN.cPPT.EF1α-TagBFP.WPRE, and the indicated viral envelope expression plasmids into HEK293T cells, followed by LVP harvest after 2–3 days. LVPs generated using the parental packaging plasmid pMDLg/pRRE served as reference controls. As negative controls, envelope-deficient LVPs were generated by replacing the envelope expression plasmid with an empty expression vector.

### LVPs generated using the new flow virometry vector system are efficiently produced, retain infectivity, and exhibit fluorescence

Lentiviruses and lentiviral vector systems may respond sensitively to genetic modifications. Indeed, insertion of the HIV-1 protease cleavage site-flanked fluorophore into HIV-1 NL4-3 to generate HIV-1 Gag-iGFP has previously been reported to reduce viral particle production, with capsid (p24) concentrations appearing to decrease by approximately half [41]. Therefore, the production efficiency of LVPs generated using the pLentiFlow vectors was assessed by quantifying p24 concentrations by ELISA and comparing them with those obtained using the parental vector system (Fig 2A).

**Fig 2.**
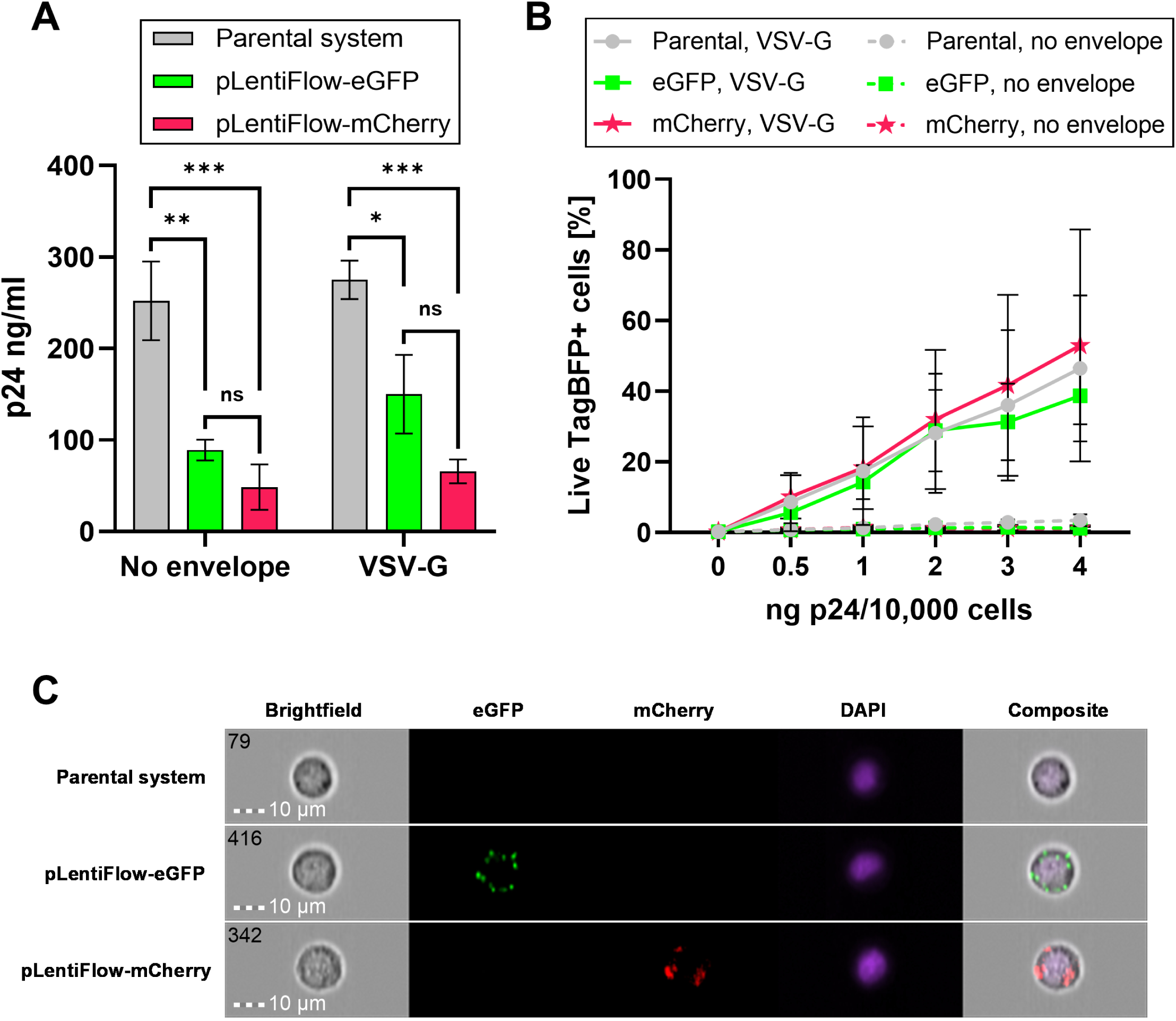
LVPs Generated Using the New Flow Virometry Vector System Are Efficiently Produced, Retain Infectivity, and Exhibit Fluorescence. **A)** Capsid protein (p24) concentrations were determined by ELISA in supernatants from HEK293T cells used to produce VSV-G-bearing or envelope-deficient LVPs with the parental or pLentiFlow-based vector systems. Bars represent the grand mean p24 concentration (ng/mL) ± standard error of the mean (SEM) from 3–6 independent LVP productions. Statistical significance was determined by two-way ANOVA with Šidák’s correction for multiple comparisons. Statistical significance is indicated as * = p < 0.05, ** = p < 0.01, *** = p < 0.001. **B)** LVPs generated using the parental system or pLentiFlow vectors and bearing VSV-G or no envelope protein were incubated with TZM-bl cells at 0–4 ng p24 per 10,000 cells. After 48–72 h, cells were harvested and infection was quantified as the percentage of TagBFP-positive cells within the live-cell population by flow cytometry. For each LVP condition, 3–4 independent LVP productions were analyzed. Data represent the grand mean ± standard deviation (SD). **C)** LVPs bearing VSV-G and generated using the parental packaging vector, pLentiFlow-eGFP, or pLentiFlow-mCherry were allowed to attach to TZM-bl cells, which were subsequently stained with DAPI and visualized by imaging flow cytometry. Representative images show brightfield, eGFP, mCherry, DAPI signal, as well as composite images.

Independent of the incorporated fluorophore and the presence or absence of an envelope protein, p24 concentrations were significantly reduced in pLentiFlow-derived LVP preparations compared with the parental system (no envelope protein: pLentiFlow-eGFP, p = 0.0011; pLentiFlow-mCherry, p = 0.0006; VSV-G: pLentiFlow-eGFP, p = 0.0141; pLentiFlow-mCherry, p = 0.0001; all compared with the parental system).

Specifically, pLentiFlow-eGFP-derived LVPs reached 35.4% (no envelope protein) and 54.6% (VSV-G) of the grand mean p24 concentrations obtained with the parental system (grand means of 252.2 and 275.1 ng p24/mL, respectively), whereas pLentiFlow-mCherry-derived LVPs reached 19.3% (no envelope protein) and 24.0% (VSV-G), respectively. Although not statistically significant, a trend towards higher particle production was observed for pLentiFlow-eGFP-derived LVPs compared with pLentiFlow-mCherry-derived LVPs both in the absence of an envelope protein (89.3 vs. 48.6 ng p24/mL, respectively) and following pseudotyping with VSV-G (150.2 vs. 65.9 ng p24/mL, respectively). Despite this reduction in particle production, pLentiFlow-derived LVPs were consistently generated at quantities sufficient for flow virometry and standard methods in HIV virology.

Next, we assessed whether modification of the Gag/Pol packaging construct affected LVP infectivity, as future applications should enable the functional characterization of bnAb-resistant particle fractions isolated by flow virometry (Fig 2B). For this, TZM-bl cells were challenged with p24-normalized amounts of LVPs generated using the parental packaging plasmid, pLentiFlow-eGFP, or pLentiFlow-mCherry. LVPs were either pseudotyped with VSV-G or bore no envelope protein. VSV-G-pseudotyped LVPs generated using pLentiFlow-eGFP or pLentiFlow-mCherry showed comparable infectivity to LVPs generated using the parental system across the tested range of 0–4 ng p24 per 10,000 cells. At the highest particle input of 4 ng p24 per 10,000 cells, the grand mean proportions of live TagBFP-positive cells were 46.5%, 38.7%, and 53.0% for LVPs generated using the parental system, pLentiFlow-eGFP, and pLentiFlow-mCherry, respectively. As expected, envelope-deficient LVPs showed only negligible transduction at the same particle input, with live TagBFP-positive cells not exceeding a grand mean of 3.49%. Together, these data demonstrate that the modifications introduced into the new flow virometry vector system do not impair LVP infectivity.

Finally, we investigated whether the incorporated fluorophores remained functional within LVPs derived from the parental or newly established vector system using imaging flow cytometry (Fig 2C). TZM-bl cells were incubated with LVPs at 4°C for 30 min to permit particle attachment while preventing viral uptake and subsequently stained with DAPI. As expected, no fluorescent particles were detected on cells incubated with LVPs generated using the parental system. In contrast, discrete green or red fluorescent signals were readily detected on cells incubated with LVPs derived from pLentiFlow-eGFP or pLentiFlow-mCherry, respectively. These findings demonstrate that the incorporated fluorescent proteins remained functional and specifically label LVPs.

Together, these results establish that the newly established lentiviral vector generates fluorescent LVPs in sufficient quantities without compromising infectivity, thereby providing a platform that is compatible with the needs for flow virometry and future functional analyses of viral subpopulations.

### VSSC-1 and an optimized threshold setting maximize detection of HIV-sized particles

The diameter of HIV-1 virions was initially reported to be approximately 120 nm by electron microscopy [42], while later studies identified a size range of approximately 110–155 nm for mature virions [43, 44]. Larger particles of up to 200 nm have also been observed, with virion immaturity or the presence of multiple cores being linked to increased particle size [43, 44]. As our analyses are primarily focused on mature virions, the relevant size range largely overlaps with the recommended operating ranges of both violet side scatter detectors of the CytoFLEX nano. According to the manufacturer, VSSC-1 is recommended for particles of approximately 40–150 nm and VSSC-2 for particles of approximately 80–1,000 nm. We therefore compared the performance of both detectors for resolving particles within the size range relevant to mature HIV-1 virions.

To identify the optimal detector settings for HIV-sized particles, we measured 100 nm polystyrene beads (Sub-micron Particle Size Reference Kit, Life Technologies) in 0.05 µm-filtered double-distilled water (ddH_2_O) while systematically titrating the avalanche photodiode (APD) gain for both VSSC channels (Fig 3A–E).

**Fig 3.**
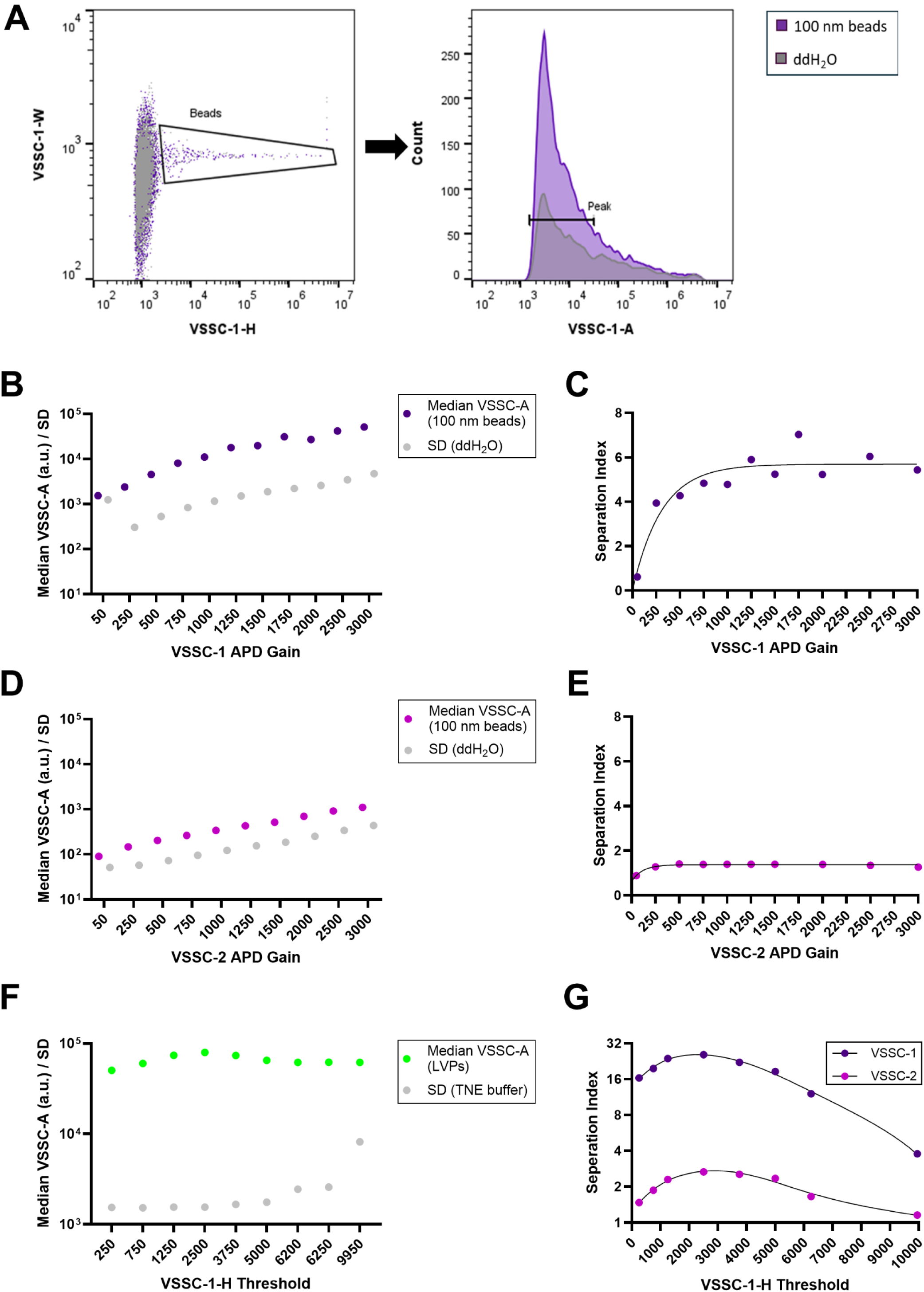
VSSC-1 and an Optimized Threshold Setting Maximize Detection of HIV-Sized Particles. **A)** Representative gating strategy for 100-nm polystyrene beads measured alongside double-distilled water (ddH₂O). The bead population was gated based on violet side scatter (VSSC)-1-H and VSSC-1-W (left), followed by identification of the peak bead population based on VSSC-1-A (right). **B, D)** Avalanche photodiode (APD) gain titrations for VSSC-1 (B) and VSSC-2 (D), showing the median VSSC-A signal of 100-nm polystyrene beads in arbitrary units (a.u.) and the standard deviation (SD) of ddH₂O measurements across the tested gain settings. **C, E)** Separation indices calculated from the measurements shown in (B) and (D) for VSSC-1 (C) and VSSC-2 (E), respectively. Curves represent non-linear regression fits. **F, G)** VSSC-1 threshold titration using pLentiFlow-eGFP-derived LVPs bearing HIV-1 NL4-3 Env. **F)** Median VSSC-1-A signals of LVPs and SDs of TNE buffer measurements across the tested VSSC-1-H threshold settings. **G)** Separation indices calculated for both VSSC-1 and VSSC-2 signals across the tested VSSC-1-H threshold settings. Curves represent smoothing splines.

Following gating of the peak bead population (Fig 3A), the median VSSC-A signal intensity of the gated bead population was determined. Background noise was quantified as the standard deviation (SD) of a Gaussian-fitted signal distribution obtained from flow cytometric measurements of ddH_2_O using the FCS Background Noise Fitting Tool [45]. Both median VSSC-A and SD were determined across the tested range of gain settings for VSSC-1 and VSSC-2 (Fig 3B, D). Median-based separation indices were subsequently calculated to determine the gain setting providing optimal signal-to-noise discrimination (Fig 3C, E) [46].

For VSSC-1 optimization, the VSSC-2 gain was kept constant at 131, while the VSSC-1 gain was titrated (Fig 3B). To maintain bead detection despite increasing background levels at higher gain settings, the VSSC-1-H threshold was adjusted from 750 to 1,000 for gain values above 1,500. Separation index analysis identified a VSSC-1 gain of 1,250 as optimal, providing robust separation of 100 nm beads from background noise (Fig 3C). For VSSC-2 optimization, the VSSC-2 gain was titrated while maintaining a constant VSSC-1-H threshold of 750 (Fig 3D). Under these conditions, a VSSC-2 gain of 500 provided stable signal-to-noise separation and was selected for subsequent analysis (Fig 3E). Comparison of the selected detector settings demonstrated superior performance of VSSC-1 for 100 nm particles, with a 4.2-fold higher separation index compared with VSSC-2 (VSSC-1: 5.89 at gain 1,250; VSSC-2: 1.40 at gain 500).

To further improve signal-to-noise discrimination, we next optimized the VSSC-1 threshold setting using fluorescent LVPs (Fig 3F, G). Since the light-scattering properties of polystyrene beads differ from those of HIV-1 particles, pLentiFlow-eGFP-derived LVPs bearing HIV-1 Env were used instead of polystyrene beads, and the SD was calculated for LVP-solvent TNE buffer (Fig 3F). Separation indices were calculated across the tested threshold range (Fig 3G). Increasing the VSSC-1 threshold improved the separation indices of both VSSC-1 and VSSC-2 in a similar manner, with the largest improvement observed at a threshold of 2,500 (maximum separation index: VSSC-1, 25.60; VSSC-2, 2.66). This threshold was therefore selected for all subsequent analyses.

Taken together, optimization of the APD gain settings demonstrated that both VSSC detectors enabled discrimination of 100 nm particles and LVPs from background noise, while VSSC-1 consistently provided superior signal-to-noise separation. Accordingly, the VSSC-1 channel was used for trigger threshold optimization, resulting in a threshold setting of 2,500 for subsequent measurements.

### Fluorescent LVPs constitute the vast majority of detected eGFP+ particles and can be reliably analyzed as single particles

Given the technical challenges associated with flow virometry, we first sought to establish that fluorescent events detected in pLentiFlow-eGFP-based LVP preparations predominantly represent LVPs rather than background or contaminating EVs. For these analyses, we first compared the ability of VSSC-1 and VSSC-2 to resolve small differences in particle size within the approximate size range of HIV-1, using NIST-traceable, size-standardized beads ranging from 80 to 200 nm (S1A Fig). VSSC-2 provided superior size resolution. In addition, arbitrary scatter and fluorescence signals were converted into estimated particle diameters and ERF units, respectively, using a refractive index of 1.42 previously determined for enveloped viruses (S1B–C Fig) [36].

Using our optimized instrument settings, a broad eGFP+ gate was applied to capture fluorescent events across a wide range of scatter-based diameters and fluorescence intensities (Fig 4A). Visualization of the detected particles revealed no obvious tail formation, characterized by an extension of the particle population toward higher scatter and fluorescence intensities and indicative of particle aggregation, despite previous reports that lentiviral aggregation increases progressively with each freeze–thaw cycle [29] and the LVPs analyzed here having undergone one such cycle. This difference may reflect variations in experimental conditions, including particle concentration or the use of TNE as the sample buffer.

**Fig 4.**
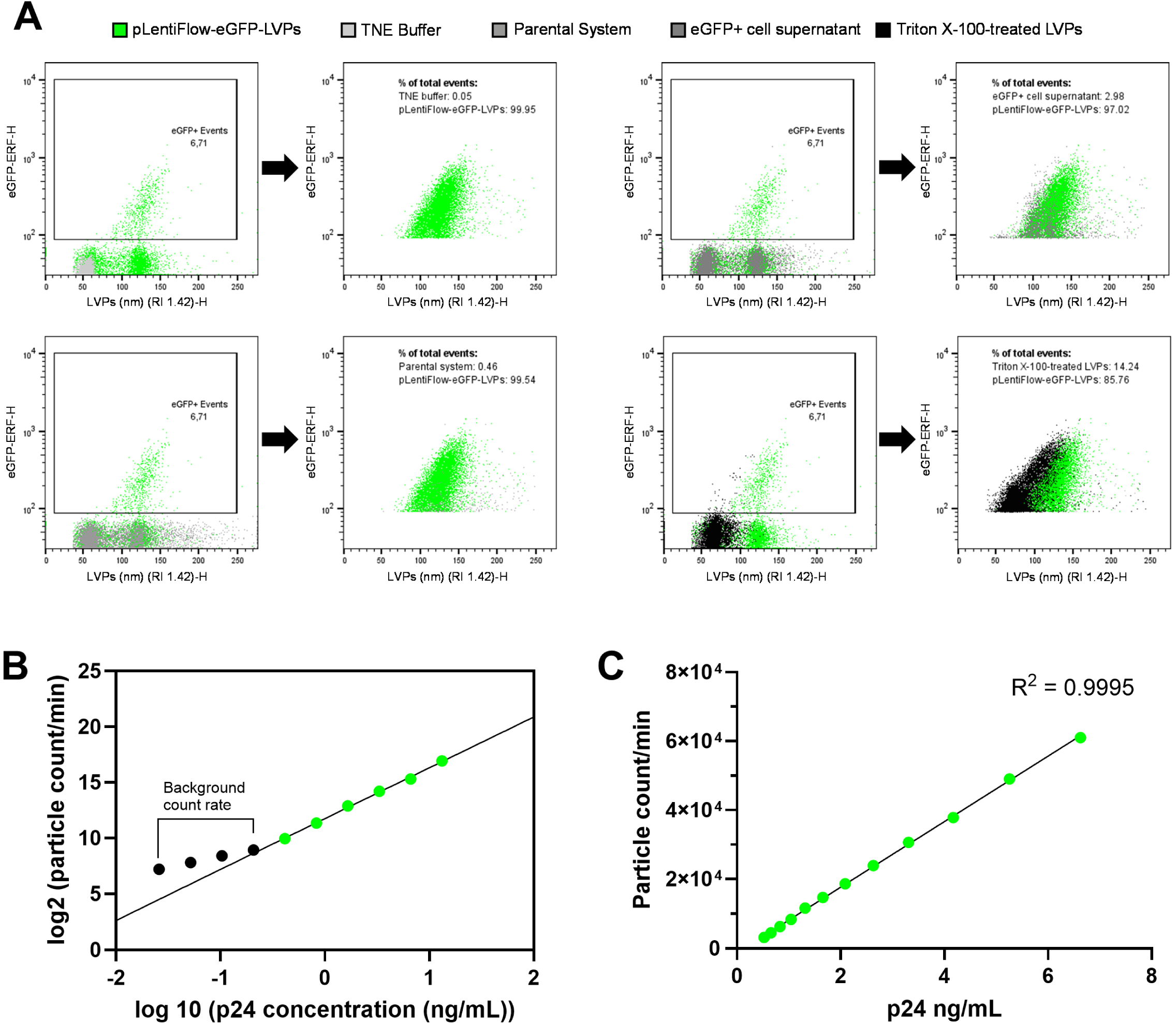
Fluorescent LVPs Constitute the Vast Majority of Detected eGFP+ Particles and Can Be Reliably Analyzed as Single Particles. **A)** Representative flow virometry plots of pLentiFlow-eGFP-derived LVPs bearing HIV-1 Env (green) overlaid with the indicated controls shown in gray or black. Controls included TNE buffer, LVPs generated using the parental system, supernatant from eGFP-transfected HEK293T cells, and Triton X-100-treated pLentiFlow-eGFP-derived LVPs. For each comparison, the left panel shows all detected events, while the right panel shows events within the eGFP+ gate. Percentages indicate the proportion of events contributed by the LVP sample and the respective control among all events within the eGFP+ gate. Scatter and fluorescence signals were converted from arbitrary units to estimated particle diameter (nm) for enveloped viruses and equivalent reference fluorophore (ERF) units, respectively. **B–C)** Serial dilutions of pLentiFlow-eGFP-derived LVPs bearing HIV-1 Env were prepared based on p24 concentration, and events within the eGFP+ gate defined in (A) were counted. **B)** LVPs were titrated from 13.25 ng p24/mL in two-fold dilution steps. Black dots represent measurements affected by instrument noise and excluded from non-linear regression analysis. **C)** Linearity was further assessed at higher resolution using a finer titration from 0.5 to 6.6 ng p24/mL, covering the sample-sparing concentration range intended for subsequent analyses. The line represents a straight-line fit generated by non-linear regression.

To determine the contribution of different sources of background within this gate, pLentiFlow-eGFP-based LVPs were measured alongside the LVP solvent, filtered TNE buffer, and non-fluorescent LVPs produced using the parental vector system (Fig 4A, upper and lower left panels). TNE buffer contributed only 0.05% of events detected within the eGFP+ gate, demonstrating minimal buffer-derived background. Similarly, samples containing non-fluorescent LVPs generated with the parental vector system, as well as EVs co-produced by the producer cells, contributed only 0.5% of events within the eGFP+ gate. Thus, noise arising from either non-fluorescent LVPs or co-produced EVs resulted in only minimal contamination of the fluorescent particle population.

However, HIV-1 Gag has been reported in particles bearing EV-associated markers, suggesting its possible incorporation into EVs [47]. Whether these particles represent bona fide EVs containing Gag remains difficult to establish, as they may instead represent Gag-driven virus-like particles budding from membrane domains enriched in EV-associated markers. Indeed, it has been demonstrated that HIV can incorporate EV-markers [48, 49]. To model the potential contribution of fluorescent EVs arising from the incorporation of eGFP-labeled Gag in the absence of LVPs, we transfected HEK293T cells with an eGFP-expressing plasmid and assessed the resulting fluorescent particles within the eGFP+ gate independently of LVP production (Fig 4A, upper right panel).

Events derived from this control accounted for 2.98% of the events detected within the eGFP+ gate. Thus, fluorescent EVs were estimated to make only a minor contribution to the detected population, supporting that the vast majority of eGFP+ events represent fluorescent LVPs.

As an additional criterion for LVP identity, we assessed whether the detected particles were membrane-enclosed. pLentiFlow-eGFP-based LVPs were treated with Triton X-100 at a concentration of 1% for 30 min (Fig 4A, lower right panel). Triton X-100 treatment markedly reduced the number of events within the eGFP+ gate, with the treated sample accounting for only 14.24% of total events instead of the 50% expected if Triton X-100 had no effect. In addition, the remaining events shifted toward smaller scatter-based diameters, consistent with partial disruption of membrane-enclosed particles. These findings further support that the detected eGFP+ events predominantly represent membrane-enclosed LVPs.

Apart from establishing the identity of the detected particles and quantifying potential background contributions, reliable single-particle analysis requires measurements to be performed within the linear concentration range of the instrument. At excessively high LVP concentrations, swarm detection can occur, in which multiple particles simultaneously passing through the interrogation volume are detected as a single event and thereby compromise single-particle analysis [50]. Conversely, at very low LVP concentrations, instrument background constitutes an increasing fraction of detected events and may therefore interfere with reliable particle detection [50]. We therefore titrated pLentiFlow-eGFP-based LVPs according to their p24 concentration and determined event counts within the eGFP+ gate (Fig 4B, C). Measurements of a two-fold dilution series starting at 13.25 ng p24/mL revealed two concentration-dependent regimes, with measurements at the lowest concentrations affected by instrument noise and a linear measurement range extending from 0.41 to 13.25 ng p24/mL that was well fitted by non-linear regression (R² = 0.9979) (Fig 4B). A subsequent finer titration from 0.5 to 6.6 ng p24/mL was performed to identify a sample-sparing concentration for subsequent measurements and confirmed linearity across this range. A straight-line fit generated by non-linear regression yielded an R² of 0.9995 (Fig 4C). Based on these results, 5 ng p24/mL was selected as the standard LVP concentration during acquisition for subsequent flow virometry measurements.

Overall, these controls demonstrate that potential background contributions to the broadly defined eGFP+ population were low, with the largest contribution arising from fluorescent EVs at approximately 3%. Detergent sensitivity further supported that the detected eGFP+ events predominantly represent membrane-enclosed LVPs. Finally, concentration-dependent titration established the linear measurement range for pLentiFlow-based LVPs on the CytoFLEX nano, enabling reliable single-particle analysis under standardized conditions.

### The flow virometry vector platform supports versatile analysis of antibody binding to diverse viral envelope proteins

Lentiviral pseudotyping enables the incorporation of heterologous envelope proteins into a common particle backbone. As a proof of concept for the application of the flow virometry vector platform, we investigated whether this versatility could be exploited to analyze antibody binding to pLentiFlow-derived LVPs bearing envelope proteins from different viruses. LVPs were generated using pLentiFlow-eGFP and pseudotyped with VSV-G, SARS-CoV-2 spike, or HIV-1 Env, while envelope-deficient LVPs served as negative controls. All tested envelope-specific antibodies were directly labeled with APC and titrated over a range of concentrations against LVPs bearing the respective envelope protein or envelope-deficient control LVPs (Fig 5). For these analyses, we focused on the core population of eGFP+ events to minimize the inclusion of aggregated particles and assess antibody binding to individual LVPs (Fig 5A).

**Fig 5.**
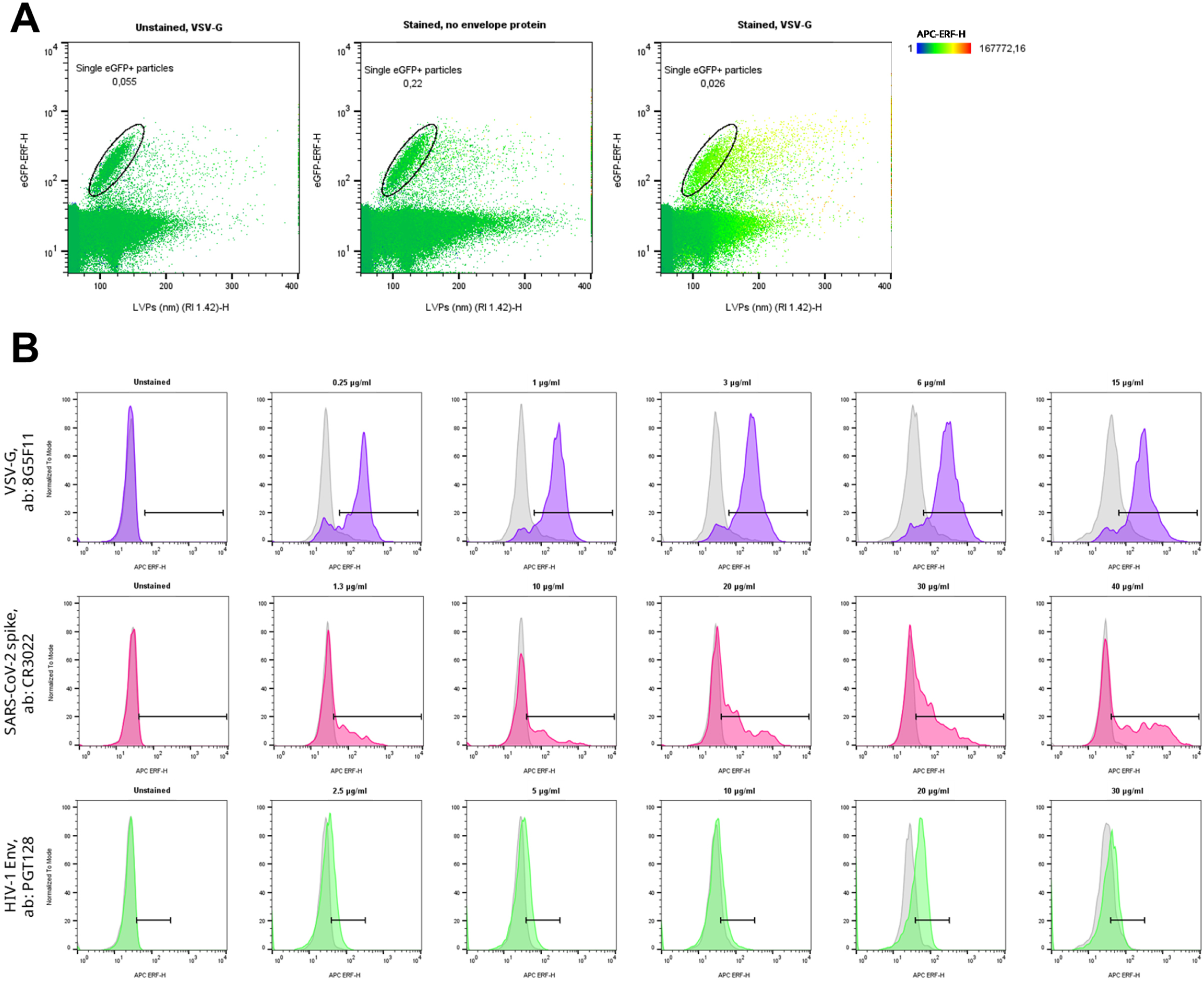
The Flow Virometry Vector Platform Supports the Versatile Analysis of Antibody Binding to Diverse Viral Envelope Proteins. **A)** Exemplary gating of the core population of eGFP+ LVPs for the subsequent assessment of individual particle antibody staining using APC-labeled antibodies. **B)** LVPs generated using pLentiFlow-eGFP and bearing either no envelope protein, VSV-G, SARS-CoV-2 spike, or HIV-1 Env were left unstained or stained with increasing concentrations of the respective envelope-specific, APC-labeled antibodies 8G5F11, CR3022, or PGT128. Histograms show the APC-ERF-H signal obtained for LVPs bearing no envelope protein (gray) or the indicated envelope protein.

For detection of VSV-G (Fig 5B, upper panel), we used the neutralizing antibody 8G5F11 (formerly I1), which binds G proteins from multiple vesiculoviruses by recognizing a conserved linear three-amino-acid epitope within the pH-sensitive domain of the G protein [51, 52]. VSV-G-bearing LVPs showed readily detectable specific staining with a clear peak separation already at an antibody concentration of 0.25 µg/mL, resulting in 71.3% specifically stained LVPs after subtraction of background staining observed for envelope-deficient control LVPs. Increasing the antibody concentration to 3 µg/mL resulted in maximal staining of 80.1% of LVPs, while background staining remained low (median APC-ERF-H: 240.0 for VSV-G-bearing LVPs vs. 29.0 for envelope-deficient LVPs). At concentrations of 6 µg/mL and above, background staining increased, reducing the proportion of specifically stained LVPs.

SARS-CoV-2 spike expression on LVPs was assessed using CR3022 [53] (Fig 5B, middle panel). CR3022 recognizes a conserved, cryptic epitope within the receptor-binding domain (RBD), distinct from the receptor-binding site, but does not neutralize SARS-CoV-2 [54–56]. Staining of SARS-CoV-2 spike-bearing LVPs with CR3022 resulted in a clearly detectable antibody-positive particle population, with 32.3% of LVPs specifically stained at an antibody concentration of 1.3 µg/mL. Specific staining increased with antibody concentration and reached a maximum of 50.2% at 30 µg/mL (median APC-ERF-H: 44.9 for SARS-CoV-2 spike-bearing LVPs vs. 26.4 for envelope-deficient LVPs). In contrast to VSV-G staining, the CR3022-positive population exhibited a broad fluorescence distribution, indicating considerable variation in staining intensity among individual antibody-positive LVPs.

Finally, HIV-1 Env was visualized using PGT128 (Fig 5B, lower panel), which recognizes an epitope within the oligomannose patch of gp120 comprising two conserved glycans (N301 and N332) and a segment of the V3 loop [57]. Despite this defined epitope, PGT128 exhibits considerable flexibility in its recognition, tolerating a two-amino-acid shift in the position of the N332 glycan-binding site or the incorporation of alternative glycans [58]. Specific PGT128 staining of HIV-1 Env-bearing LVPs was first detectable at an antibody concentration of 2.5 µg/mL, resulting in 25.5% specifically stained LVPs, and reached a maximum of 51.2% at 20 µg/mL (median APC-ERF-H: 48.4 for HIV-1 Env-bearing LVPs vs. 26.7 for envelope-deficient LVPs). Compared with VSV-G, the antibody-positive population showed less pronounced separation from the envelope-deficient control population.

Together, these data demonstrate that envelope proteins from three different viruses can be specifically detected on individual pLentiFlow-derived LVPs, demonstrating that the pseudotyping capacity of the lentiviral system can be exploited to analyze different viral envelope proteins while maintaining an otherwise consistent LVP composition.

Across the analyzed envelope proteins, specific antibody staining was detected on a substantial proportion, but not all, of the LVP population. Specific staining could be resolved above background, although with less pronounced signal-to-background separation than typically observed in conventional cellular flow cytometry, consistent with the limited epitope numbers expected on small viral particles. Moreover, these titrations demonstrated that the antibody concentration required for maximal specific staining varied between LVPs bearing different envelope proteins, emphasizing the importance of optimizing staining conditions for maximal envelope detection.

## Discussion

In this study, we developed a third-generation lentiviral vector system for flow virometry. By transferring the design principle of HIV Gag-iGFP to a third-generation lentiviral vector backbone, we generated two fluorescent packaging constructs carrying either eGFP or mCherry. Similar to the previously described HIV Gag-iGFP reporter virus [41], insertion of fluorophores into the Gag polyprotein resulted in reduced capsid production while preserving infectivity comparable to the parental system. Importantly, the LVPs produced with the new flow virometry vector system were fluorescently detectable, enabling their visualization and discrimination from similarly sized sample and buffer contaminants. Together with the retained infectivity of these particles, their intrinsic fluorescence may facilitate downstream applications requiring the identification and functional characterization of defined LVP subpopulations. We further optimized measurement conditions, establishing VSSC-1 as the optimal parameter for event triggering and thresholding and VSSC-2 for scatter-based particle analysis. Using a variety of controls, we confirmed that pLentiFlow-eGFP-derived LVPs are membrane-enclosed particles and that background from the assessed sources accounted for no more than ∼3% of eGFP+ events, indicating that the vast majority represent fluorescent LVPs. We further defined a linear measurement range. Finally, as a proof of concept, we demonstrated the versatility of the flow virometry system for analyzing antibody binding to LVPs bearing diverse viral envelope proteins, including VSV-G, SARS-CoV-2 spike, and HIV-1 Env.

Across all three envelope proteins, antibody staining identified a substantial but incomplete fraction of the fluorescent LVP population, with the highest proportion of specifically stained particles observed for VSV-G, reaching approximately 80% at optimal antibody concentrations. Several factors may contribute to these differences. First, although our controls indicated that background from the assessed sources accounted for no more than ∼3% of eGFP+ events, the contribution of fluorescent EVs may have been underestimated because free eGFP used in the EV control may be incorporated into EVs less efficiently than Gag-associated eGFP. Consequently, a proportion of the analyzed eGFP+ events may represent fluorescent EVs that recruited fluorescent Gag but did not carry viral envelope proteins.

In addition, differences in envelope protein incorporation may contribute to the proportion of antibody-positive LVPs. Authentic virions have been reported to contain envelope protein trimer numbers of approximately 400 VSV-G [59], 24 ± 9 SARS-CoV-2 spike [60], and only 8–10 HIV-1 Env [61]. Although these numbers may differ in LVPs, they illustrate the substantial differences in envelope abundance among the three viral proteins. Moreover, HIV-1 Env and SARS-CoV-2 spike incorporation varies considerably between individual virions [60–62], raising the possibility that some LVPs carried no or little envelope protein and therefore remained below the limit of antibody detection.

The antibody used and accessibility of its epitope are also likely to influence the proportion of stained particles. Indeed, previous flow virometry screening of a large panel of HIV-1 Env-specific antibodies demonstrated substantial differences in antibody performance, with 36% of the 85 antibodies tested not yielding detectable staining above background [63]. The VSV-G-specific antibody used in this work recognizes a conserved linear epitope [51, 52], and may therefore recognize the majority of incorporated VSV-G molecules. In contrast, CR3022 recognizes a cryptic epitope on SARS-CoV-2 spike [54, 55]. Its accessibility is thought to require two RBDs in the up position, a conformation observed in only approximately 14% of the pooled prefusion spike-trimer population on virions, as well as additional, so far not naturally observed structural rearrangements that further open the spike trimer [55, 60]. PGT128 used here for visualization of HIV-1 Env recognizes an epitope on gp120 comprising a short segment of the V3 region and two glycans within the oligomannose patch [57].

Interestingly, PGT128 exhibits considerable flexibility in its recognition, tolerating a two-amino-acid shift in the position of the N332 glycan-binding site or the incorporation of alternative glycans [58]. Nevertheless, particle-to-particle variation in local gp120 conformation, glycan-site occupancy, or glycan species composition may still leave some Env-bearing LVPs undetected.

Target destabilization may further contribute to incomplete staining. For HIV-1, gp120 shedding from Env may occur spontaneously and varies among viral strains [64]. It may also be promoted by LVP processing or handling. Notably, NL4-3 Env is susceptible to cold-induced shedding, which could contribute to gp120 loss during low-temperature steps done in preparation for flow virometry [65]. Similarly, CR3022 Fab binding has been demonstrated to destabilize and disrupt the prefusion spike trimer into protomers, potentially facilitating release of the RBD-containing S1 subunit together with bound CR3022 from cleaved spike [66, 67]. Thus, the different proportions of antibody-positive LVPs observed here may reflect a combination of fluorescent EVs within the analyzed population, differences in envelope abundance and particle-to-particle incorporation, antibody-dependent epitope recognition, and target loss. Together, these considerations highlight how little is currently known about the envelope protein profiles of individual particles within viral populations and about how variations in these profiles may influence antibody recognition.

Addressing these questions will likely require complementary experimental approaches. While our platform enables controlled investigations of the contribution of Env to viral heterogeneity, alternative approaches are more suitable when the focus is the direct analysis of donor-derived HIV populations. For example, bead-based capture has been used to isolate HIV particles directly from donor plasma, including virus derived from macrophages through CD36 capture and from CD4+ T cells through CD27 capture [68]. Using this approach, the authors demonstrated that macrophage-derived, but not T cell-derived, virus carries bioactive transforming growth factor beta on its surface, suggesting a potential role for producer cell-dependent viral phenotypes in HIV pathogenesis. While this approach yielded interesting insights, bead coupling may associate multiple viral particles with a single bead and has so far not been demonstrated to be reversible for downstream infection experiments.

In contrast, direct antibody labeling of host cell markers enabled the sorting of infectious HIV from in vitro-produced virus samples, while Env-specific labeling allowed the isolation of individual bnAb-reactive HIV particles for quasispecies analysis [69]. These findings demonstrate the versatility of direct antibody labeling. However, EVs can carry host cell markers used for HIV isolation, including CD44 and CD36 [70, 71], and may in some cases also incorporate HIV proteins such as Env and Gag [47, 72, 73].

Consequently, isolation strategies based on a single target molecule may not sufficiently discriminate between HIV particles and EVs, potentially contributing to the low recovery rate of approximately 0.5% reported for the isolation of simian/human immunodeficiency virus from plasma [69].

Together, both studies highlight the growing potential of flow virometry for the investigation of HIV heterogeneity directly in donor-derived samples. At the same time, approaches based on donor-derived viral populations and controlled pseudotyping systems should be viewed as complementary rather than competing technologies. While direct analyses provide access to naturally occurring viral populations, they offer only limited opportunities to dissect the contribution of individual viral features to the observed phenotypes. HIV-1 Env represents a particularly relevant source of non-genetic viral variation, as individual particles can differ in Env glycosylation, stability, and conformation [15–18]. Once viral variants of interest have been identified and sequenced from donor samples, pseudotyping approaches such as the system presented here could allow the contribution of Env to be investigated in isolation on an otherwise identical viral backbone.

In summary, our third-generation lentiviral vector system for flow virometry provides a versatile framework for single-particle analysis of antibody binding to viral envelope proteins. Combined with complementary approaches for the direct analysis of naturally occurring viral populations, this system may contribute to a better understanding of envelope-associated non-genetic diversity in HIV and other viruses and the consequences of such variation for antibody recognition. Such insights may advance research on antibody-based vaccines and therapeutics.

## Acknowledgements

We acknowledge that these experiments utilized reagents (p24-ELISA components) provided by the AIDS and Cancer Virus Program, Biological Products Core Laboratory, Frederick National Laboratory for Cancer Research, supported with federal funds from the National Cancer Institute, National Institutes of Health, under contract 75N91019D00024. The following reagents were obtained through BEI Resources, NIAID, NIH: TZM-bl cells (HRP-8129; contributors: J.C. Kappes, X. Wu, and Tranzyme, Inc.); expression plasmids encoding VSV-G (ARP-4693; contributor: Lung-Ji Chang), SARS-CoV-2 Wuhan-Hu-1 spike (NR-52310; contributors: F. Krammer, F. Amanat, and S. Strohmeier; produced under HHSN272201400008C), HIV-1 NL4-3 Env (ARP-324; contributors: E. Freed and R. Risser, [74]), and the heavy- and light-chain expression plasmids for the anti-SARS coronavirus monoclonal antibody CR3022 (NR-53260; contributors: F. Krammer, V. Chromikova, and G. Asthagiri Arunkumar; produced under HHSN272201400008C); the antibody PGT128 (ARP-13352; contributor: IAVI; [57, 75]) against HIV-1 gp120. Undermind was used to complement classical literature research and support literature organization. Language editing was performed by ChatGPT (OpenAI). All scientific content was written and validated by the authors.

