## Supplementary material for "A Third-generation Lentiviral Vector System for Flow Virometry-based Analysis of HIV-1 Env and Other Viral Envelope Proteins": S1 Fig

### Supporting information

**S1 Fig. VSSC-2 Resolves Small Particle Size Differences in the HIV Size Range and Enables Scatter-Based Size Estimation of LVPs. A)** Polystyrene beads with diameters of 80, 125, and 200 nm diluted in $ddH_2O$ were measured on the CytoFLEX nano using VSSC-1 (left) and VSSC-2 (right). Histograms show the scatter signal distributions of the three bead populations. **B)** Median and coefficient of variation of VSSC-2 median scatter height signals of the 80-, 125-, and 200-nm polystyrene beads shown in (A) were used to generate a scatter-diameter calibration curve using $FCM_{PASS}$. A refractive index of 1.42 was applied for enveloped viruses to derive the corresponding diameter calibration for LVPs. Red dots indicate the measured calibration bead data. **C)** Representative $FCM_{PASS}$ model-fit output generated as part of the scatter-diameter calibration shown in B), displaying the agreement between measured calibration bead scatter signals and model predictions. Abbreviations: Acq., acquired; predict., predicted, RI, refractive index.

### S1 Fig

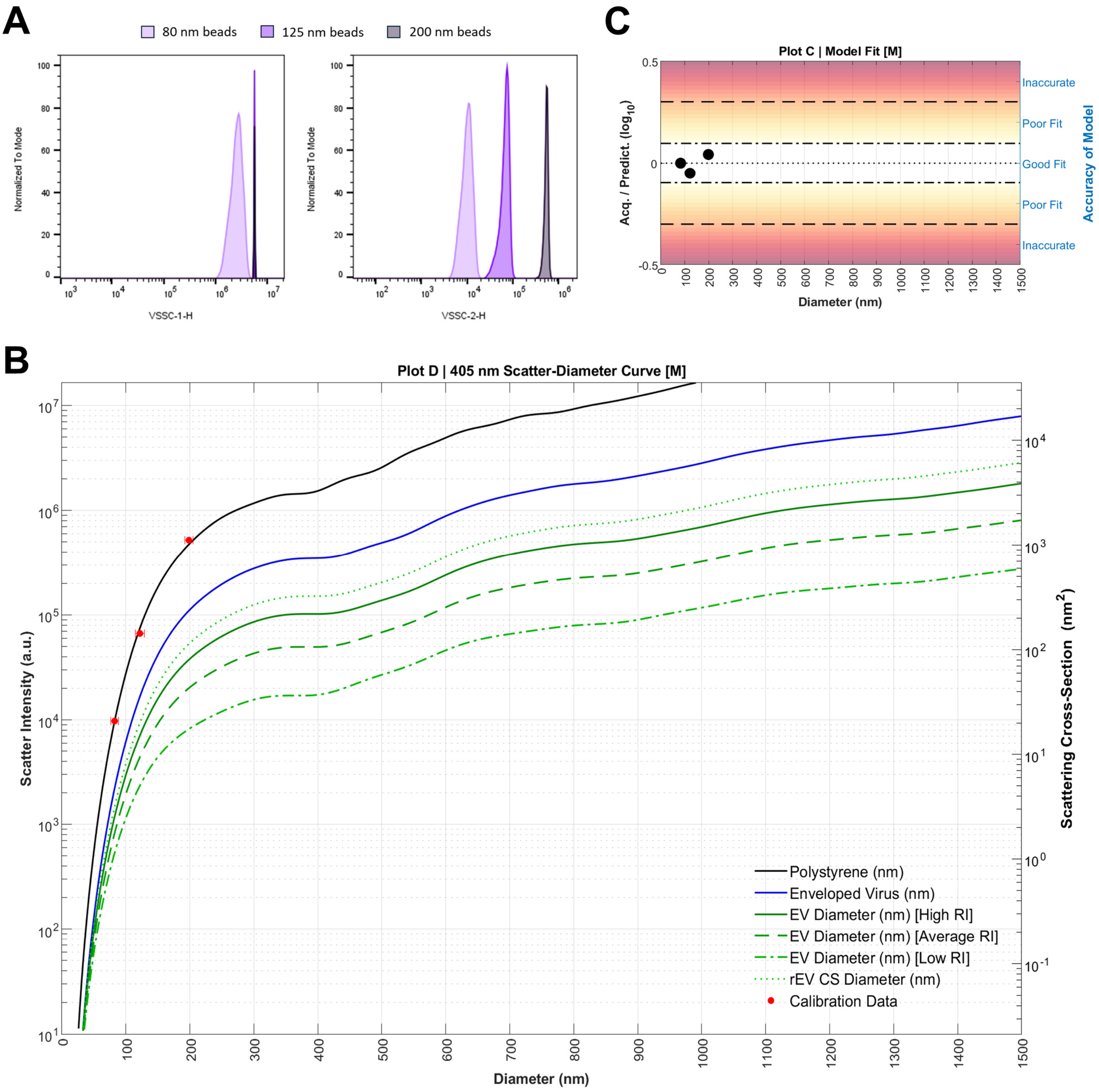

A
80 nm beads
125 nm beads
200 nm beads
Normalized To Mode
VSSC-1-H
VSSC-2-H
C
Plot C | Model Fit [M]
Acq. / Predict. (log10)
Inaccurate
Poor Fit
Good Fit
Poor Fit
Inaccurate
Accuracy of Model
Diameter (nm)
B
Plot D | 405 nm Scatter-Diameter Curve [M]
Scatter Intensity (a.u.)
Scattering Cross-Section (nm2)
Diameter (nm)
Polystyrene (nm)
Enveloped Virus (nm)
EV Diameter (nm) [High RI]
EV Diameter (nm) [Average RI]
EV Diameter (nm) [Low RI]
rEV CS Diameter (nm)
Calibration Data
